# A Proteomic Platform to Identify Off-Target Proteins Associated with Therapeutic Modalities that Induce Protein Degradation or Gene Silencing

**DOI:** 10.1101/2020.11.18.389148

**Authors:** Xin Liu, Ye Zhang, Lucas D. Ward, Qinghong Yan, Tanggis Bohnuud, Rocio Hernandez, Socheata Lao, Jing Yuan, Fan Fan

## Abstract

Novel modalities such as Proteolysis Targeting Chimera (PROTAC) and RNA interference (RNAi) have a mechanism of action-based potential to alter the abundance of off-target proteins. The current *in vitro* secondary pharmacology assays, which evaluate off-target binding or activity of small molecules, do not fully assess the off-target effects of PROTAC and are not applicable to RNAi. To address this gap, we developed a proteomics-based platform to comprehensively evaluated abundance of off-target proteins. The first part of the manuscript describes the rationale and process through which the off-target proteins and cell lines were selected. The off-target proteins were selected from the entire human proteome based on genetics and pharmacology data (Deaton *et al*., 2018). The selection yielded 2,813 proteins, forming the nexus of a panel that we refer to as the “selected off-target proteome” (SOTP). An algorithm was then used to identify appropriate cell lines. Four human cell lines out of 932 were selected that, collectively, expressed ~ 80% of the SOTP based on transcriptome data. The second part of the manuscript describes the LC-MS/MS experimentation to quantify the intracellular and extracellular proteins of interest in the 4 selected cell lines. Among over 10,000 quantifiable proteins identified, 1,828 were part of the predefined SOTP. The SOTP was designed to be easily modified or expanded, owing rationale selection process developed and the label free LC-MS/MS approach chosen. This versatility inherent to our platform is essential to design fit-for-purpose studies that can address the dynamic questions faced in investigative toxicology.

## INTRODUCTION

Adverse drug reactions (ADRs) are one of the main contributors to drug attrition during clinical development and post-marketing drug withdrawal (Roberts et al., 2014; Hamon et al.). Hence, an effective ADR assessment during nonclinical development is beneficial to both patients and the pharmaceutical industry. Roughly 75% of ADRs are associated with the pharmacological profile of candidate compounds (Bowes et al., 2012), which can be further separated into primary (on-target) and secondary (off-target) effects (Smith and Schmid, 2006; Redfern et al., 2002). Secondary pharmacology screening is often performed by pharmaceutical companies as a cost-effective approach to assess unexpected binding and/or activity against a panel of common off-target proteins; however, existing in vitro assays are mainly applicable for small molecules or peptides. For emerging therapeutic modalities such as Proteolysis Targeting Chimera (PROTAC) and RNA interference (RNAi), the assessment of potential secondary pharmacology effects requires new testing methods. A PROTAC is a hetero-bifunctional molecule that is composed of two ligands, one binding to a protein of interest and the other recruiting an E3 ubiquitin ligase, connected by a linker. PROTACs achieve target degradation via the proteasome mediated ubiquitination machinery (Pettersson and Crews, 2019). With the continued exploration of therapeutic targets into the realm of “undruggable” proteins, (Hopkins and Groom, 2002), there has been increasing interest in developing PROTAC as a new therapeutic modality. RNAi is a gene therapy approach that is intended to silence targeted genes, via RNA-induced silencing complex (RISC). RNAi has emerged from a research tool to a therapeutic modality and has moved rapidly into the clinical trials. Both modalities have the potential for off-target activity, which would be reflected by changes in protein abundance, rather than binding or activity modulation.

The protein level changes that can result from off-target activity of a PROTAC or a RNAi molecule involve key cellular components including the ubiquitin-proteasome system and RISC, respectively. Thus such off-target activities can only be assessed within the correct biological context and by using appropriate test systems. In fact, the importance of preserving biological relevance in off-target identification was demonstrated by a study using a small molecule β-secretase inhibitor (Zuhl *et al*., 2016), for which cathepsin D was identified as the key driver of off-target activity using living cells rather than in vitro biochemical assays. Although such detailed case studies have yet to be performed with PROTAC or RNAi largely due to their early stages as therapeutics, the use of living cells provides the necessary biological context to properly evaluate protein degradation or knockdown.

To this end, we have developed a novel proteome-based platform in human cell lines to support appropriate off-target evaluation of modalities such as PROTAC and RNAi. We systematically selected off-target proteins from the entire human proteome based on phenotypes from genetic and pharmacological evidence, in a similar fashion to previous work from our group (Deaton *et al*., 2018), including proteins that are implicated in the function of the cardiovascular, respiratory and central nervous systems. These 2,813 proteins form the panel that we refer to as the “selected off-target proteome” (SOTP). Instead of making recombinant protein or engineered overexpression cell lines for each off-target in the SOTP, our cell-based platform uses native endogenous proteins for the screening assay. The transcriptomics data from 932 cell lines were used to identify expression of genes in the SOTP and the whole genome. Using a Greedy algorithm, 4 cell lines were selected with maximized gene transcription coverage for both the SOTP genes and the whole protein-coding transcriptome. Global proteomics and detailed characterization were then used to identify the quantifiable proteins from both cytosolic and secreted fractions. Using the intensity-based absolute quantification, the expression levels of proteins in the SOTP were found to differ by up to six orders of magnitude, suggesting that the system could detect the change of protein abundance even across a wide range of expression. Despite the selection focused only on 3 key organ systems our analysis showed the SOTP, at its current composition, offers good coverage in all key organs, defined in the MedDRA hierarchy at the highest level (system organ classes), allowing the platform to be used to probe off-target activities while focusing on certain organ systems of interests. The conservation of the amino acid sequence of the 2,813 proteins across human, rat, dog, and monkey were also calculated and suggest that this platform approach could be useful to help guide the selection of relevant animal species to test off-target activity. In summary, the proteome-based platform we developed is a valuable tool to assess relevant off-target effects of new modalities such as PROTAC and RNAi.

### MATERIALS AND METHODS

#### Database Selection and SOTP Identification

The pharmacology and genetics databases used in this study were previously compiled (Nguyen *et al*., 2019). Briefly, drugs and their intended targets for generating the pharmacology database were comprehensively obtained from DrugBank (Wishart *et al*., 2018), Citeline Pharmaprojects (Pharma Intelligence, Informa PLC., d2016-11-22), and a recently curated database of therapeutic efficacy targets of a subset of marketed drugs (Santos *et al*., 2019). Intended targets were obtained from the union of the databases and target annotations were cross referenced between the databases. Our database of pharmacological evidence was built by pairing the targets and the indications of these drugs.

Human genes and corresponding phenotypes were derived from the Human Phenotype Ontology (HPO), STOPGAP, and GWAS Catalog databases. For associations from the GWAS Catalog, only associations with a genome-wide significant p-value < 5E-8 were used for subsequent analysis. Our database of genetic evidence was generated by linking genes to the phenotypes from these databases.

To enable aggregation of phenotypes across databases, the Unified Medical Language System (UMLS) Metathesaurus was used to map phenotypic terms and ADRs in the previous described pharmacology and genetics databases using the MetaMap natural language processing (NLP) tool, and the UMLS-Interface software. More specifically, phenotypic terms and ADR terms were mapped to Medical Dictionary for Regulatory Activities terminology (MedDRA) standard and only the system organ classes (SOC) terms were retained for downstream analysis. Thereafter, drug targets and genes that involved in heart, vascular, nervous, respiratory, and mental SOC terms were extracted from the databases to identify the 2,813 genes that comprise the SOTP.

#### Comparison of SOTP between Human and Nonclinical Species

The protein sequences of the SOTP and their corresponding orthologs in *R.norvegicus* (rat), *C. familiaris* (dog), and *M. fascicularis* (macaque), were obtained from the OMA orthology database using OMA IDs as queries (Altenhoff *et al*., 2018). These data were downloaded from the PyOMADB library (Kaleb *et al*., 2019) using a python script based on OMA Browser’s REST API. With the sequence pairs downloaded, pairwise global Needleman-Wunsch sequence alignments were conducted with BioPython (Cock *et al*., 2009). The similarity scores were then calculated by dividing the number of identical amino acids by the total length of the alignment between two species. The result ranges from 0 to 1, with 0 annotating the ortholog as not found and 1 indicating an identical alignment. The resulting similarity score list only includes the human proteins with at least one ortholog identified from the OMA orthology database. The similarity scores for SOTP were visualized with Seaborn boxplot method (Waskom *et al*., 2017). In addition, we extracted class and cellular localization information from IPA and provided this information together with the similarity scores (Qiagen; Krämer *et al*., 2014).

#### Cell Line Selection

Gene expression data of 932 cancer cell lines from the Broad Institute’s Cancer Cell Line Encyclopedia (CCLE, version 2012-Sept) RNA sequencing dataset, which contains the RNA expression data for 932 cancer cell lines, was analyzed for the study (Barretina et al., 2012). The analysis was carried out using OmicSoft’s Array Studio software (www.omicsoft.com/array-studio.php; Oshell.exe v9.0). The expression values were quantile normalized at the 70th percentile to value of 10 (FPKQ, fragments per kilobase per million reads; quantile normalized). We define genes with FPKQ >= 1 as being expressed in the corresponding cell line.

Then, we ordered and selected an experimentally manageable set of cell lines (i.e., ideally 3-5) to collectively express a considerable portion of the proteome. Here, we implemented a greedy algorithm to identify an approximation of the set of cell lines that achieve an optimal coverage of the proteome. (Supplementary Figure 11) Briefly, the algorithm iteratively selects the cell line that would result in the best cumulative coverage either over the SOTP or over whole proteome. After iterating over all tested cell lines, the algorithm generates an ordered list of cell lines, from which the top 3 cell lines with the maximal cover for SOTP and whole proteome were selected.

#### Cell Culture

Human medullary thyroid carcinoma (MTC) cell line TT and human pancreatic cancer cell line SU.86.86 were purchased from the American Type Culture Collection (ATCC). Human esophageal squamous cell carcinoma cell line KYSE-270 and Human small cell lung carcinoma (SCLC) cell line COR-L24 were obtained from European Collection of Authenticated Cell Cultures (ECACC). TT cells were cultured in ATCC-formulated F-12K medium supplemented with 10% fetal bovine serum (FBS) (Hyclone, UT, USA). SU.86.86 cells were grown in RPMI 1640 medium containing 10% FBS. KYSE-270 cells were maintained in RPMI 1640 and Ham’s F12 (Invitrogen, CA, USA) mixed (1:1) medium containing 2mM L-glutamine (Invitrogen, CA, USA) and 2% FBS. COR-L24 cells grew in aggregates and were cultured in RPMI 1640 medium supplemented with 2mM L-glutamine and 10% FBS. All cell lines were maintained at 37 °C in humidified air containing 5% CO_2_.

#### Cellular Proteome Sample Preparation

For mass spectrometric analysis, the schematic workflow is shown in Figure 1 Intracellular proteins (referred to as the cellular proteome) were collected by harvesting three consecutive passages of cells as biological triplicates. Adherent cells (TT, SU.86.86, and KYSE-270) were washed three times with ice-cold phosphate-buffered saline (PBS) (Invitrogen, CA, USA) and then harvested by scraping into another 1 mL of ice-cold PBS before centrifugation at 1,000 × g for 5 min at 4 °C. COR-L24 suspension cells were first centrifuged at 300 × g for 5 min to pellet the cells and then washed three times with ice-cold PBS. The supernatants were discarded, and the adherent and suspension cell pellets were solubilized in lysis buffer containing 5% sodium dodecylsulfate (SDS) (Sigma-Aldrich, MO, USA) in 50 mM triethylammonium bicarbonate (TEAB, pH 7.6) (Sigma-Aldrich, MO, USA) at room temperature (RT). A high concentration of SDS detergent was also utilized to help dissolve and increase coverage of poorly-soluble membrane proteins. To shear the DNA and reduce the lysate’s viscosity, the samples were sonicated on ice for three 30 s rounds at 25 % amplitude with a probe sonicator (Agilent, CA, USA). SDS lysates were heated to 90 °C for 10 min and clarified at 15, 000 × g for 10 min. Total protein concentrations were determined using the bicinchoninic acid (BCA) protein assay kit (Thermo Fisher Scientific, IL, USA). Total amounts of 600 μg proteins were then reduced with 20 mM dithiothreitol (DTT) (Sigma-Aldrich, MO, USA) for 30 min at 56 °C and then alkylated with 40 mM iodoacetamide (IAA) (Thermo Scientific Pierce, MA, USA) for 30 min in the dark. After quenching with an additional of 20 mM DTT at RT for 30 min, protein digestion in the S-Trap filter was performed according to manufacturer’s instructions with slight modifications. Briefly, to the sample was added a final concentration of 1.2% phosphoric acid and then six volumes of binding buffer (90 % methanol; 100 mM TEAB, pH 7.1). After gentle mixing, the protein solutions were loaded to S-Trap filters, spun at 4,000 × g for 30s, and the flow-throughs collected were reloaded onto the filters. This step was repeated twice, and then the filters were washed three times with binding buffer. Finally, trypsin was added at 1:20 (wt:wt) in 50 mM TEAB (pH 8), and digested overnight at 37 °C. To elute peptides, three step-wise buffers were applied, with 50 mM TEAB, 0.2% formic acid (FA) (Thermo Scientific Pierce, MA, USA), and 60% acetonitrile and 0.2% FA. All eluents containing tryptic peptides were pooled together and vacuum-centrifuged to dryness.

**Figure 1.**
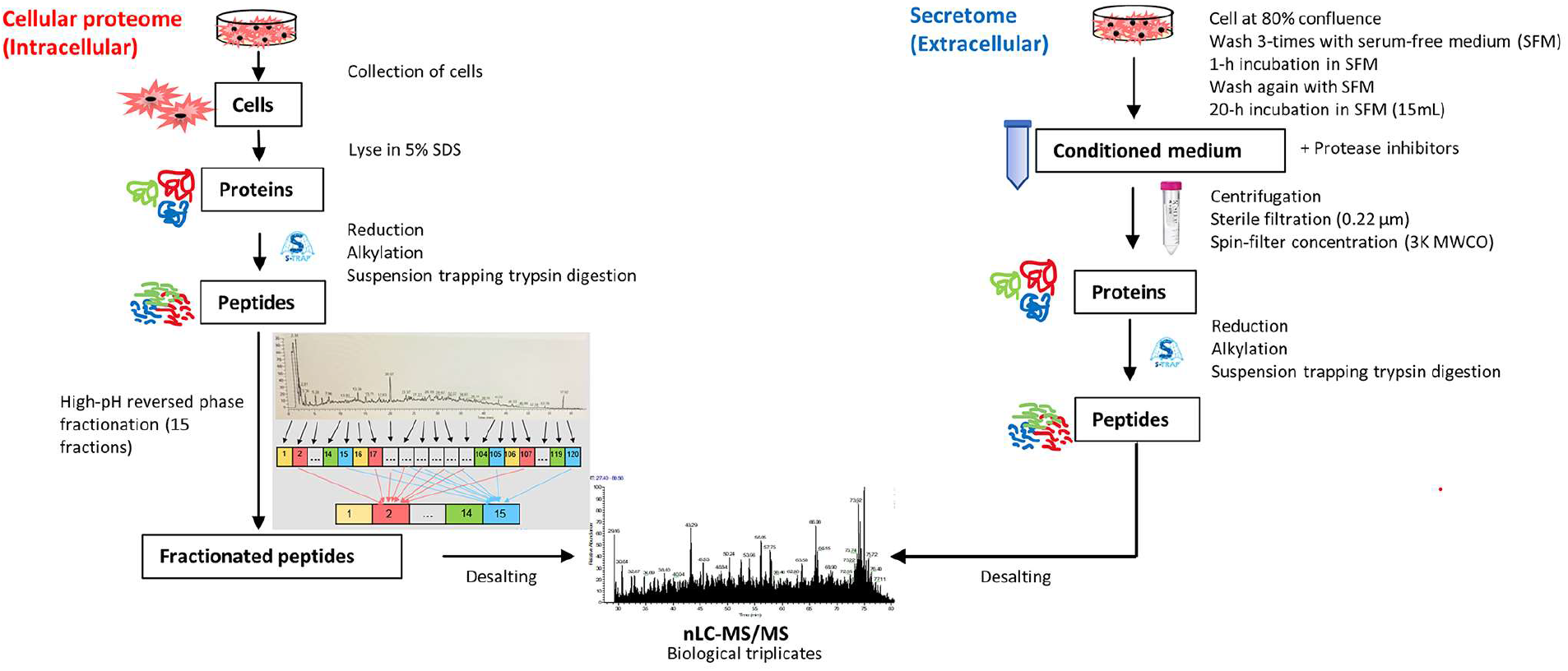
Schematic workflow for the quantification of cellular proteome and secretome.

Dried peptides were resuspended in 15 mM ammonium bicarbonate (ABC) (Sigma-Aldrich, MO, USA) and fractionated using a Waters XBridge BEH130 C18 3.5 μm 4.6 × 150 mm column on a Vanquish UHPLC system (Thermo Fisher Scientific, NY, USA) operating at 1 mL/min with buffer A consisted of 15 mM ABC at pH 8 and buffer B consisted of 15 mM ABC with 95% acetonitrile, pH 8. Peptides were separated by a linear gradient from 2% B to 35% B in 50 min followed by a linear increase to 60% B in 7 min, and ramped to 70%B in 3 min. At this point, fraction collection was halted, and the gradient was increased to 98% B in 9 min before being ramped back to 2% B, where the column was then washed and equilibrated. Fractions were collected at 30s intervals to a total of 120 fractions and were then recombined by pooling every 15^th^ fraction in a step-wise concatenation strategy to yield a total of 15 fractions. All fractions were dried by vacuum centrifugation, resuspended in 0.1% FA in water and desalted. For nanoflow LC-MS/MS, the loading amount was kept constant at 1 μg per injection, determined by quantitative colorimetric peptide assay (Thermo Fisher Scientific, IL, USA).

#### Secretome Sample Preparation

To collect the secretomes, adherent cells at 80% of confluence in 75 cm^2^ flasks were washed three times with sterile PBS, while COL-R24 suspension cells were centrifuged first and then rinsed with PBS. The cells were then exposed to serum free medium (SFM) for 1 h, rinsed once again with SFM and incubated with 15 mL of SFM for 20 h. After incubation, the cell viability was monitored with trypan blue dye exclusion to be over 90% for each cell line. 1% (v/v) EDTA-free protease inhibitor cocktail (Roche, IN, USA) was added to the collected cell supernatants (referred to as the secretome). The conditioned media were then centrifuged at 3000 × g for 15 min at 4 °C, and sterile-filtered through a 0.22 μm filter unit (Millipore, MA, USA) to remove cell debris. The supernatants were then concentrated and desalted with water via Amicon 3 kDa filter device (Millipore, MA, USA) at 4 °C, and protein concentrations were determined by BCA assay. 30 μg of proteins were reduced, alkylated, and digested the same way using S-Trap filters as described above. Digested peptides were kept at −80 °C before LC-MS/MS analysis.

#### Liquid Chromatography and Mass Spectrometry Analysis

Digested samples were analyzed using a Q-Exactive hybrid quadrupole-Orbitrap mass spectrometer (Thermo Fisher Scientific, MA, USA) coupled to an Ultimate 3000 RSLCnano system (Thermo Fisher Scientific, MA, USA) through an EASY-Spray ion source (Thermo Fisher Scientific, MA, USA). Chromatographic separation of the peptides was performed on an EASY-Spray C18 column (75 cm × 75 μm inner diameter, packed with PepMap RSLC C18 material, 2 μm) at a flow rate of 0.25 μL/min. Solvent A consisted of 0.1% formic acid (FA) in water, while solvent B consisted of 0.1% FA in acetonitrile (ACN). The following gradient was used for all samples:2% B for 0-5 min, 2-30% B from 5-110 min, 30-55% B from 110-130 min, 55-90% B from 130-140 min, 90% B until 155 min, and re-equilibration at 2% B from 155-180 min. All solvents were liquid chromatography mass spectrometry grade. The mass spectrometer was operated in Top 12 data-dependent mode with automated switching between MS and MS/MS. Capillary temperature was maintained at 300 °C and the ion source was operated in positive ion mode at 2.0 kV. Full MS scans were acquired from 380 to 1800 m/z at a resolution of 70, 000, with an AGC target of 1 × 10^6^ ions and a fill time of 230 ms. MS^2^ scans were performed from 100 to 1500 m/z at a resolution of 17, 500 and a maximum fill time of 120 ms. The AGC target was set at 1 × 10^5^ ions with an underfill ratio of 0.4%. An isolation window of 1.4 m/z was used for fragmentation with a normalized collision energy of 30. Dynamic exclusion was set at 40 s. Ions with a charge of +1 or greater than +7 were excluded from fragmentation.

#### Computational Mass Spectrometric Data Analysis

Raw MS files were analyzed by MaxQuant software (version 1.6.4.0) equipped with the Andromeda search engine. MS/MS spectra were searched against the Uniprot human database (20,416 sequences) concatenated with 248 common contaminants. For secretome MS data search, a list of FBS associated proteins was also included (Shin *et al*., 2019). A first search was performed with a precursor mass tolerance of 20 ppm, the results of which were used for mass recalibration. In the main search, precursor mass and product ion mass had an initial mass tolerance of 4.5 ppm and 20 ppm, respectively. Trypsin was set as the digestion enzyme with a maximum of two missed cleavages and minimal peptide length was set to six amino acids. Carbamidomethylation was set as a fixed modification, while oxidation (M), acetylation (protein N-term), and deamidation (NQ) were set as variable modifications. Target decoy analysis was performed by searching a reverse database with an overall false discovery rate (FDR) of 0.01 for peptide and protein identifications. For label-free protein quantification, the XIC-based MaxLFQ was used. The algorithm first calculated pairwise protein ratios by taking the median of all pairwise peptide ratios per protein to protect against outliers. Only shared identical peptides were considered for each pairwise comparison with a minimum of one ratio count. The relative abundance profile for each protein was then reconstructed with a least-squares analysis. To maximize the number of quantification events across biological samples within each cell line, we enabled the “match between runs” feature with a matching time window of 0.7 min and an alignment time window of 20 min to allow the quantification of high-resolution MS1 features that were not identified in each single measurement. For estimation of the absolute abundance of different proteins within a single sample, we used the intensity-based absolute quantification (iBAQ) algorithm. The values are the intensities divided by the number of theoretical peptides. Thus, iBAQ levels are proportional to the molar quantities of the proteins. Lysates and secretomes were analyzed as two independent batches. The data output from Maxquant was analyzed using Perseus software (version 1.6.5.0), R or Python frameworks.

Proteins that were marked as contaminants, identified only by site modification or found in the decoy reverse database, were excluded. For quantitative analysis, LFQ intensities (normalized intensities) were log2 transformed and only proteins with at least one identified unique peptide and a minimum of two valid values in at least one cell line were considered. Missing data were imputed by values from a normal distribution (width 0.3 standard deviations) of down-shifted 1.8 standard deviations. Hierarchical clustering of proteins was performed after z-score normalization of the data, using Euclidean algorithm with Ward’s linkage method. Principal Component Analysis (PCA) of cell lines relied on singular value decomposition and the original feature (protein) space was orthogonally transformed into a set of linearly uncorrelated variables (principal components). These account for distinct types of variation in the data. For pairwise comparison of proteomes, a two-sided t-test was used with a S0 constant of 2 and a permutation-based FDR of 0.05. Presented fold changes have been calculated as difference from mean values of log2 transformed intensities. Multiple t-tests (ANOVA) was performed with FDR value of 0.01. Cellular compartment data and protein classes were obtained from Uniprot, Ingenuity Pathway Analysis (IPA), PANTHER Classification System data analysis tool (version 14.1), or DAVID Bioinformatics Resources (version 6.8).

For secretome analysis, proteins were classified using bioinformatics databases. The classically secreted proteins were searched using “Signal” or “Secreted” as keywords in Uniprot, or were identified using Signal Peptide Predictor (SignalP, version 5.0). SignalP uses amino acid sequences to predict the presence of signal peptides and cleavage sites with a probability score of ≥ 0.9. To identify nonclassical, or leaderless, protein secretion, SecretomeP (version 2.0) was used. SecretomeP is a neural network-based method that has used six protein features to determine if a protein is non-classically secreted. These characteristics include: number of atoms, number of positively charged residues, presence of transmembrane helices, presence of low-complexity regions, pro-peptide cleavage site, and subcellular localization. A protein is considered non-classically secreted if it receives an NNscore of ≥ 0.5. Moreover, it is also possible that proteins located on the plasma membrane are shed and released to the extracellular space. Therefore, TMHMM (version 2.0) was used to predict transmembrane helices. Finally, the exosome proteins were matched by the ExoCarta database cause such proteins may not pass the SignalP and SecretomeP score cut-offs.

For functional class evaluation, a list of 1,158 mitochondrial genes was obtained from MitoCarta 2.0 (Calvo *et al*., 2016). Drug target genes obtained from Drugbank (v.5.0.6) and restricted to proteins related to MOA for at least one drug. Transcription factor (TF) genes (n=1,639) were from the Human TFs collection (Lambert *et al*., 2018). Disease associated genes were acquired from Uniprot, which is a knowledgebase consisting of both manually annotated records with information from literature and information from OMIM database. Cancer-related genes, including mutated and cancer driver genes across 21 tumor types as well as genes implicated in malignant transformation, were downloaded from COSMIC (Futreal *et al*., 2004). For analysis of ubiquitination essential genes, the list of 929 ubiquitination (UBQ)-related genes including E1, E2 enzymes as well as E3 ligases and their associated adaptor genes and 95 deubiquitinating genes were obtained (Ge *et al*., 2018).

### RESULTS

Systematic selection of the SOTP. To address the need for comprehensive off-target profiling especially for new drug modalities, we aimed to establish a comprehensive cell-based, proteomics-centered platform. One way to build such a platform is to achieve high coverage of the whole human proteome, but this would be costly both to develop and use. Therefore, we prioritized a subset of proteins to include in this pilot study by focusing on targets involved in major organ systems including cardiovascular, respiratory, and central nervous systems, as highlighted by ICH-S7A (ICH, 2000), following a method previously outlined by our group (Deaton et al., 2018). To expand our previous work, we constructed a database of SNPs, genes and annotations pertaining to large scale GWAS studies, Mendelian traits, drug adverse effect, and drug indications (Deaton et al., 2018). We performed a series of phenotypic mappings and queries to gather genetic and pharmacological evidence for targets that are implicated in the aforementioned organ systems. Genetic evidence alone identified 2,423 proteins to include in the SOTP (Figure 2A), while pharmacology evidence contributed 248 proteins. A total number of 142 proteins were identified by both genetic and pharmacology data.

**Figure 2.**
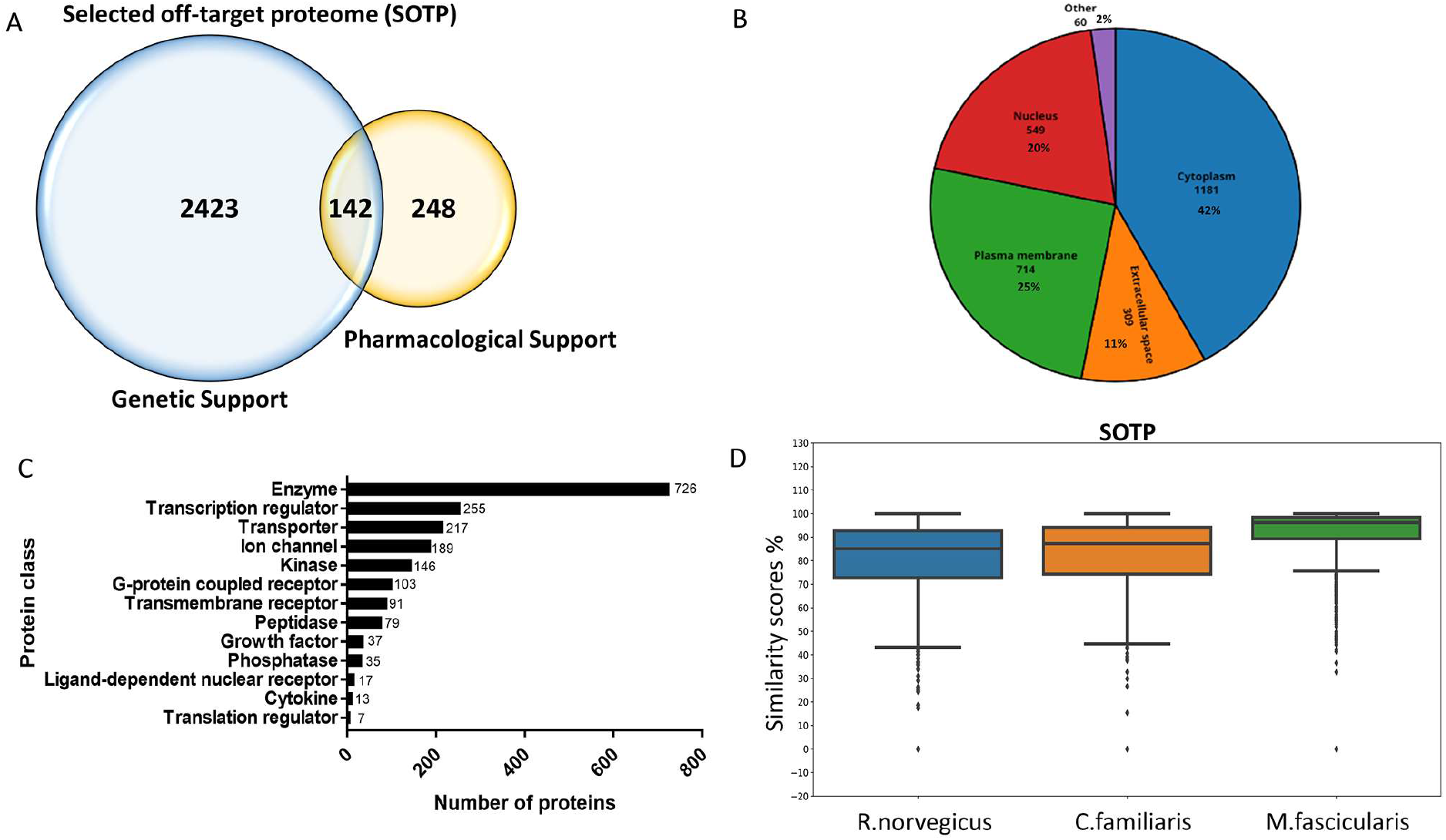
Systematic selection of SOTP and cell lines. (A) Venn diagram of SOTP composition. (B) Subcellular localization of SOTP. (C) Protein class of SOTP. (D) the similarity of primary protein structures between species.

The 2,813 selected proteins were analyzed in detail for various attributes, including subcellular localization, target classes, the organ systems they belong to, as well as similarity of amino acid sequences across species. The subcellular locations of the selected proteins were mainly distributed between the cytoplasm (42%), plasma membrane (25%), nucleus (20%), and extracellular space (11%) (Figure 2B). Based on Uniprot classification of each protein in the SOTP, the largest target classes include enzymes, transcription regulators, transporters and ion channels as shown in Figure 2C. The organ systems where each protein is involved was also analyzed, based on evidence from genetics, pharmacology and biological pathways. The populated implications were then mapped to the highest MedDRA level, i.e., system organ class. A good coverage, i.e., an average of 67%, was observed across all the key MedDRA systems, such as liver, kidney, gastrointestinal, eye, skin and so on. A complete list of the 2,813 proteins, their subcellular location, target class, as well as the organ systems they are involved in, are detailed in the Supplementary Data (Supplementary Table 1).

Secondary pharmacology screening mainly focuses on human proteins. However, it is often important to assess the potential for off-target activity across nonclinical species to help predict potential translatability of findings observed in nonclinical studies or identify a relevant model to further understand findings observed in the clinic. To this end, we compared the amino acid sequence for each human protein in the SOTP with that of its corresponding orthologs in three nonclinical species, namely *R. norvegicus, C. familiaris*, and *M. fascicularis*. The similarity scores range from 0 to 1. A similarity score of 1 means that the protein sequences are identical. As expected, the results indicated a higher protein sequence similarity between human and macaques than human and rat or dog (Figure 2D). The similarity scores for the 2,813 proteins in the SOTP were listed in the Supplementary Table 1.

Systematic selection of cell lines. To enable good coverage of the SOTP with a manageable number of cell lines, we utilized transcriptomic data from 932 cell lines (Barretina et al., 2012). Maximizing the combined transcriptomic coverage of multiple cell lines would ideally be calculated through an exhaustive search algorithm of all possible combinations, which would be computationally prohibitive. However, we hypothesized that iterative addition of lines with the highest SOTP coverage rank would rapidly reach a coverage plateau since the total number of proteins in our signature is only 2,813. We identified four cell lines whose transcriptome data suggested that they express greater than 80% of the SOTP (more than 2,000 proteins). With the expanding body of genetic and pharmacological knowledge over time, a broader panel of safety-related proteins may be established in the future. Therefore, to generalize our platform, we also attempted optimizing the coverage of the whole human proteome. Utilizing the same approach, genes expressed by top three ranked cell lines, SU8686, COR-L24, and TT, cover 73% of the whole proteome. Collectively, to allow both manageable experimentation and decent target coverage of both the SOTP and the whole proteome, four cell lines were selected: SU8686, COR-L24, KYS-270, and TT. COR-L24 and KYS-270, from male donors, were lung cancer and esophagus cancer cell lines, respectively. SU8686 and TT, from female donors, were pancreatic cancer and thyroid cancer cell lines.

Quantitative proteomic profiling of selected cell lines. To make our platform expandable for modified application in future studies, we used label-free quantification with no limits on the number of samples to be analyzed for the relative quantification of proteins across cell lines. Collectively, the combined analysis of triplicates of the four selected cell lines using peptide and protein FDR thresholds of 1%, quantified protein groups (proteins distinguishable by MS) corresponded to 10,627 ENSEMBL genes, which covers 53% of the protein-coding human genome and 65% of the SOTP (Figure 3A). We required proteins to be quantified in at least two biological replicates of at least one cell line (≥1 unique peptide). Only 3% of the quantified proteome was quantified with 1 unique peptide. The genes encoding identified proteins were evenly distributed across chromosomes (Supplementary Figure 2).

**Figure 3.**
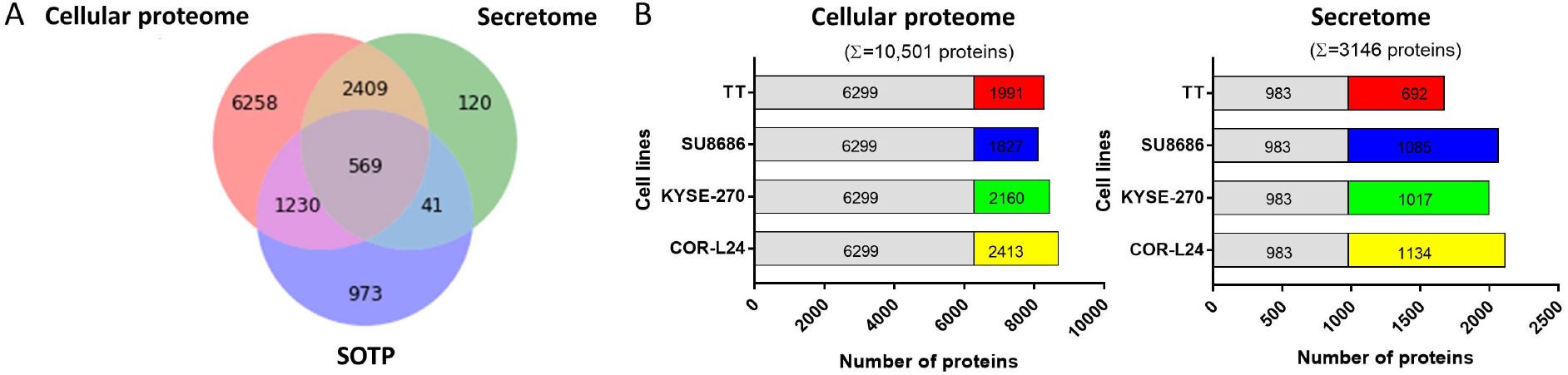
Deep proteomic analysis of selected cell lines. (A) Venn diagram of proteins quantified in cellular proteome and secretome compared to SOTP (gene centric). (B) Proportion of proteins quantified in all and contributions of different cell lines.

The proteomic profiling consists of two separate experiments, namely cellular proteome analysis and secretome analysis. For cellular proteome analysis, intracellular proteins were extracted from cells and 10,501 unique protein groups were quantified with 173,705 unique peptides (Figure 3B and Table 1). The median number of unique tryptic peptides per protein was 11, leading to an average sequence coverage of 36%. When the cell lines were analyzed separately, 8,000-9,000 proteins were quantified in each of them. A total number of 6,299 proteins were quantified ubiquitously in all four cell lines, and the remaining ~4,000 proteins show a more distinct expression pattern with ~2,000 proteins contributed by each cell line. For secretome analysis, secreted proteins were collected from conditioned medium and 3,146 protein groups were quantified with 35,230 unique peptides (Figure 3B and Table 1). The median number of unique tryptic peptides per protein was 7, leading to an average sequence coverage of 29%. In each cell line, approximately 2,000 proteins were quantified. A total number of 983 proteins were quantified ubiquitously in all four cell lines with approximately 1,000 proteins contributed by each cell line. Among the quantified proteins, 694 (22.1 %) proteins were identified as classical secreted proteins marked with the keywords “Signal” or “Secreted” in UniProtKB or predicted by SignalP containing a signal peptide (Supplementary Figure S3). Apart from the classical secreted proteins, 986 were predicted as nonclassical secreted proteins by SecretomeP, 122 were predicted to be integral membrane proteins, and 983 were matched by the ExoCarta exosome database. These extracellular proteins are secreted by cells through nonclassical or exosome-mediated secretion pathways, and they are vital components of the cell secretome. Collectively, these proteins accounted for 88.7% of all quantified proteins in the secretome.

**Table 1.** Protein quantification in 4 cell lines. Summary of the number of quantified proteins from biological triplicate analysis of each cell line in cellular proteome or secretome.

| | Cell lines | Quantified proteins (gene centric, 1% FDR) | # of cell-type enriched proteins (gene centric, ≥ 2FC, 1% FDR) | Peptides (unique, 1% FDR) | Peptides /protein (unique, median) | Sequence coverage (%)/protein (unique, mean) | Total quantified proteins (gene centric, 1% FDR) |
| --- | --- | --- | --- | --- | --- | --- | --- |
| **Cellular proteome** | COR-L24 | 8712 | 819 | 173705 | 11 | 36 | 10466 |
|  | KYSE-270 | 8459 | 249 |  |  |  |  |
|  | SU8686 | 8126 | 311 |  |  |  |  |
|  | TT | 8290 | 526 |  |  |  |  |
| **Secretome** | COR-L24 | 2117 | 242 | 35230 | 7 | 29 | 3139 |
|  | KYSE-270 | 2000 | 193 |  |  |  |  |
|  | SU8686 | 2068 | 294 |  |  |  |  |
|  | TT | 1675 | 242 |  |  |  |  |

To compare protein levels across the various cell lines, MS signals of the same peptides detected in different cell lines are compared to each other. Protein abundance distributions of all 4 cell lines are generally very similar (Supplementary Figure S4). To further estimate the relative abundance of proteins within a proteome, the MS signals of the peptides identifying a protein are summed and normalized to the number of theoretically observable peptides of the protein. In each of the 4 cell lines, the iBAQ values varied over above six orders of magnitude in the cellular proteome (Supplementary Figure 5) and above five orders of magnitude in the secretome (Supplementary Figure 6). The median iBAQ values across the cell lines and the estimated absolute abundance of quantified proteins of the composite cell lines proteome showed similar dynamic range of protein expression like the individual proteomes (Supplementary Figure 7). These observations are consistent with other studies estimating protein abundances in mammalian cell lines (Gholami et al., 2013; Katsogiannou et al., 2019; Coscia et al., 2016). This broad dynamic range allows for proteome-wide unbiased detection of drug off-target liabilities. Reproducibility of the label-free protein quantification between biological replicates, similarities and dissimilarities of cell lines on a global scale were then evaluated as shown in Supplementary Figures 8&9. Utilizing these cell lines with differentially expressed proteins will effectively expand the coverage and reduce false negative and false positive results.

Protein cell line expression distributions were also mirrored by functional categories of genes (Figure 4A). For example, 88% of mitochondrial genes were quantified in 4 cell lines. Among these quantified proteins, 86% were found across all cell lines indicating their central roles for maintaining cellular homeostasis. In contrast, the expression distribution of proteins classified as therapeutic targets and TFs was much more cell type restricted with only ~40% quantified in all cell lines. This is consistent with the notion that proteins may make for better drug targets if they are not ubiquitously expressed (Hao and Tatonetti, 2016). TFs are also known to be very divergently expressed related to the functional specialization of different cell types. Apart from these, the relative distribution of disease-associated genes and cancer-related genes followed that of all quantified genes. To support the use this platform for PROTAC off-target identification, we also checked for coverage of ubiquitination (UBQ)-related genes and determined that 72% of these UBQ essential genes were detected in the chosen cell lines (Figure 4A).

**Figure 4.**
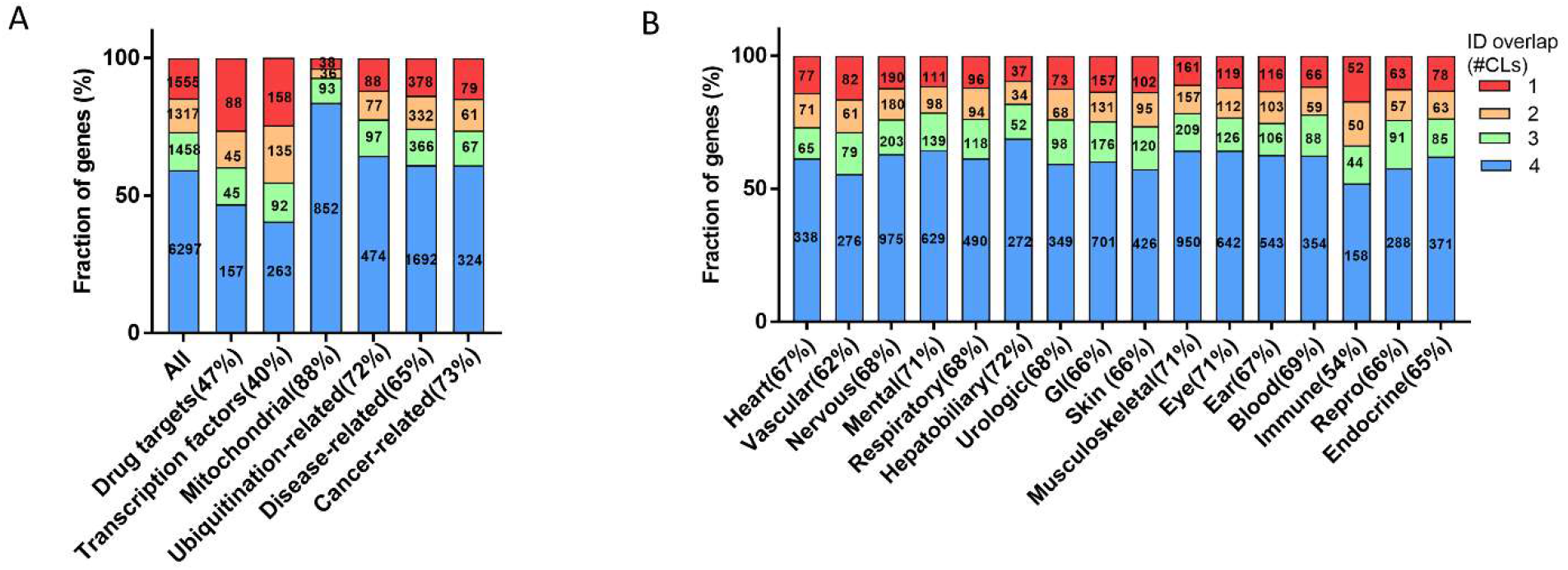
Relative distribution and absolute numbers of proteins in selected functional categories (A) and different organ classes (B). The coloring of the bars indicates fractions of proteins that are expressed in all or detected in certain number of cell lines.

Next, we analyzed the coverage and relative distribution of quantified proteins in different organ classes (Figure 4B). The lists of targets in multiple organ classes that could manifest into safety crucial phenotypes were obtained using the same approach as generating the SOTP (Supplementary Figure 10). An average coverage of 67% was achieved across different organ classes. The good coverage across all key MedDRA organs demonstrates the advantage of a large and carefully curated panel of off-target proteins.

## DISCUSSION

Unintended off-target activity is a hurdle in drug discovery not only for small molecule drug candidates, but also for emerging modalities such as PROTAC (Bondeson et al., 2018) and RNAi (Jackson and Linsley, 2010). The current approach of in vitro secondary pharmacology screening does not detect protein level changes and, therefore, does not fully characterize the potential off-target effects for these novel modalities. As both RNAi and PROTAC are relatively new drug modalities, approaches for conducting effective off-target screening for these modalities remains to be clearly defined. To address this gap, we developed a platform using living human cells that natively express a subset of proteins that are of interest from a safety standpoint, which we termed as SOTP. In addition, the SOTP platform also affords other applications, such as to address the translatability of off-target activities across species, to assess the existence of polypharmacology, as well as to facilitate possible drug repurposing.

### The SOTP platform addresses the unmet need for off-target assessment of new modalities by enabling the monitoring of protein level changes

Potential off-target activities associated with traditional small molecules are typically limited to binding, inhibition or activation, which are often dissociable and low in affinity. However, PROTAC and RNAi molecules may result in unexpected protein level changes of off-target proteins, presenting a different mechanism of action for potential off-target effects. The SOTP platform was designed to properly and systematically assess these “new” types of off-target activities for these emerging therapeutic agents.

As PROTAC is still in early an emerging modality for therapeutic drugs, there is no well characterized off-target activity assessment platforms for this modality. There are two types of off-target activities of a PROTAC molecule: small-molecule like off-target activities (typically binding, inhibition or activation) and off-target PROTAC activity, leading to the degradation of unintended target(s). Unexpected degradation is most directly reflected by the changes in protein abundance and would not be detected by current off-target assays that evaluate in vitro binding or activity. The SOTP platform provides an opportunity to systematically and comprehensively address the capability of PROTACs to degrade off-targets, which may lead to toxic phenotypes.

The SOTP platform could also be leveraged to evaluate off-target activity for another emerging modality, RNAi. The promiscuity of RNAi was demonstrated to be a prevalent issue in a detailed study performed by Lin et al. In this study, 5 out of 6 targets that were previously reported to be essential for the proliferation of cancer cells were knocked out with no impact on the survival of cancer cell lines. Such misidentification was attributed to off-target effects of the RNAi used during the initial characterization of these targets (Lin et al., 2019). One source of RNAi off-target activity comes from the partial sequence complementation between the three prime untranslated region (3’ UTR) of the off target transcripts and the 5’ end of the transfected RNAi guide strand (Jackson and Linsley, 2010). An off-target effect can result with sequence complementarity of as little as 8 nucleotides (Jackson and Linsley, 2010), which may lead to unanticipated phenotypical consequences. Currently, the off-target activity of RNAi is typically assessed by genome scale mRNA expression analysis. However, the changes in mRNA level do not often manifest into changes in protein level due to reasons such as translation rate, protein half-life, protein synthesis delay and so on (Liu et al., 2016). It is therefore important to monitor the changes in protein level upon the treatment of RNAi for a more robust off-target assessment.

The utility of the SOTP platform is not limited to emerging drug modalities as it could also be used to support conventional small molecule drug candidates. Small molecule drugs acting through a covalent mechanism can easily be detected using mass spectrometry, because the covalently linked complex does not dissociate during *the gas-phase used for* detection. The resulting drug-small molecule complex can be subsequently attached with a tag such as biotin, via bioconjugation reaction namely click chemistry (Kolb et al., 2001), which enables high affinity purification from the a pool of homogenized tissues or organs for accurate off-target identification, as exemplified by the study with inhibitors of the T790M mutant form of EGFR (Niessen et al., 2017). For non-covalently acting small molecule drugs, photo reactive modification allows an otherwise noncovalent binder to form covalent bonds with the protein backbones when UV light is applied. There has been increasing success at identifying off-targets in tissue or cell using photoaffinity labeling, as elegantly demonstrated in the mechanistic elucidation of retinal toxicity caused by β-secretase inhibitor (Zuhl et al., 2016).

As large molecules are not typically challenged with selectivity issues, the SOTP platform is not a priori intended to support these modalities; however, the platform could be adapted to support large molecule drug development. In order to achieve this, additional methods for co-immunoprecipitation and detection of cell membrane proteins would need to be developed, as large molecules such as antibodies will only interact with extracellular instead of intracellular proteins.

### Other potential applications of SOTP platform

The SOTP provides biological relevance, as the selected human off-target proteins are presented within their native cellular context. As a result, the utility of the SOTP platform expands beyond a comprehensive screen for off-target proteins.

First, during nonclinical development the potential translatability of an off-target effect across species often needs to be evaluated. This is commonly addressed using in vitro assays to compare off-target activities of proteins from human and nonclinical species. However, assays are often not readily available for all protein orthologs across multiple species and hence would require resources for reagent generation (e.g., recombinant proteins or cell lines) as well as assay development. In contrast, using the computational approaches outlined in our study, cell lines from relevant nonclinical species could be selected that express a large number of off-target proteins. Thus the off-target activities can be assessed using the same global proteomics approach utilized in human cell lines.

Second, the SOTP can be easily adapted to illustrate the polypharmacology profile of a drug molecule. Polypharmacology refers to the activities of one drug molecule against multiple targets (Peters, Jens-Uwe, 2013; Poornima et al., 2016; Anighoro et al., 2014; Hopkins, 2008; Tan et al., 2016; Hu and Bajorath, 2013). Polypharmacology drugs can be more effective for complex systematic diseases such as cancer, cardiovascular and psychiatric diseases (Poornima et al., 2016; Peters, Jens-Uwe, 2013). Similarly, adverse phenotypes also often result from the action of multiple off-target proteins. Hence, if activities were observed against many related off-targets that are linked to one adverse event, a comprehensive panel may better predict the possible phenotypical consequences. For example, the Drug Abuse Potential Profiling panel offered by Eurofins, which contains in vitro binding assays for 44 targets, represents one such effort (Eurofins discovery). Our SOTP assays a larger number of targets in living cells. The collective actions of related off-targets might paint a clearer picture to forecast possible phenotypic outcome(s).

Last, the SOTP platform may also enable repurposing an existing drug. As elegantly highlighted by Lin et al., drugs may achieve efficacy via off-target(s) rather than the intended target (Lin et al., 2019). The SOTP, with its large target coverage and biological relevance, is an ideal platform to identify additional targets that are modulated by the molecule, which may also result in better understanding of therapeutic mechanism of action, and afford the possibility of efficiently predicting, or even mitigating, target-induced toxicity.

**In summary**, as a new screening paradigm, our proteomics-based platform utilizes human cell lines as a display library to allow unbiased and biologically relevant screening for off-target proteins. As an example, we focused on key safety phenotypes and systematically selected 2,813 proteins. The SOTP platform is especially suited to support RNAi and PROTAC, for which the existing in vitro assays are not well suited for off-target identification. The SOTP platform can be used for extensive screening as well as for retrospective issue resolution. We intend to continuously update the list of off-target proteins included in the SOTP as well as the cell lines being used by repeating the computational approaches outlined in this manuscript in order to reflect the increasing knowledge of human proteins as well as to make the system most appropriate for different issue resolution situations. To this end, our SOTP platform is highly customizable, as the label free detection allows easy addition of new cell lines on to the panel used in our pilot study. To our knowledge, our platform offers the largest target coverage for off-target screening efforts and can also be easily adapted for other applications, such as drug repurposing and polypharmacology characterization. Taken together, our platform represents a step to realize the vision of early safety evaluation, where the aim is to predict potential adverse events from the molecular mechanism of toxicity, especially for new modalities.

## Supporting information

supporting information, full table

## ABBREVIATIONS

ADR: adverse drug reaction;
RNAi: RNA interference;
small interfering RNAs: siRNAs;
PROTAC: Proteolysis Targeting Chimera;
SOTP: selected off-target proteome;
MedDRA: Medical Dictionary for Regulatory Activities;
SOC: system organ classes;
FDR: false discovery rate.

## SUPPLEMENTARY DATA

Supplementary data are available at Toxicological Sciences online.

## ACKNOWLEDGMENT

The authors thank Victor Cee (Medicinal Chemistry at Amgen) as well as Herve Lebrec and Edward K. Lobenhofer (Translational Safety and Bioanalytical Sciences at Amgen) for their valuable discussion and insights.

