## Supplementary material for "A Proteomic Platform to Identify Off-Target Proteins Associated with Therapeutic Modalities that Induce Protein Degradation or Gene Silencing": figures of supporting information

**Supplementary Figure 1**

Venn diagram safety proteome composition in CV, CNS, and respiratory (gene centric).

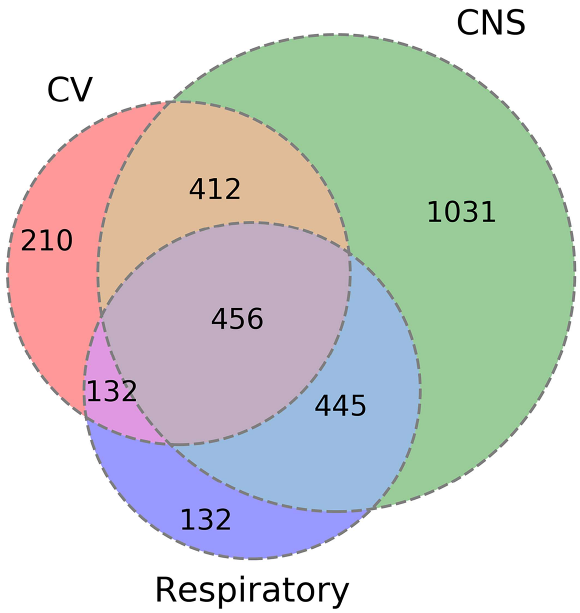

**Supplementary Figure 2**
Coverage of the human genome by chromosomes. Distributions of the quantified proteins (gene centric) are shown in red (cellular proteins) and green (secreted proteins), safety proteome are shown in orange versus total genes (blue) for each chromosome.

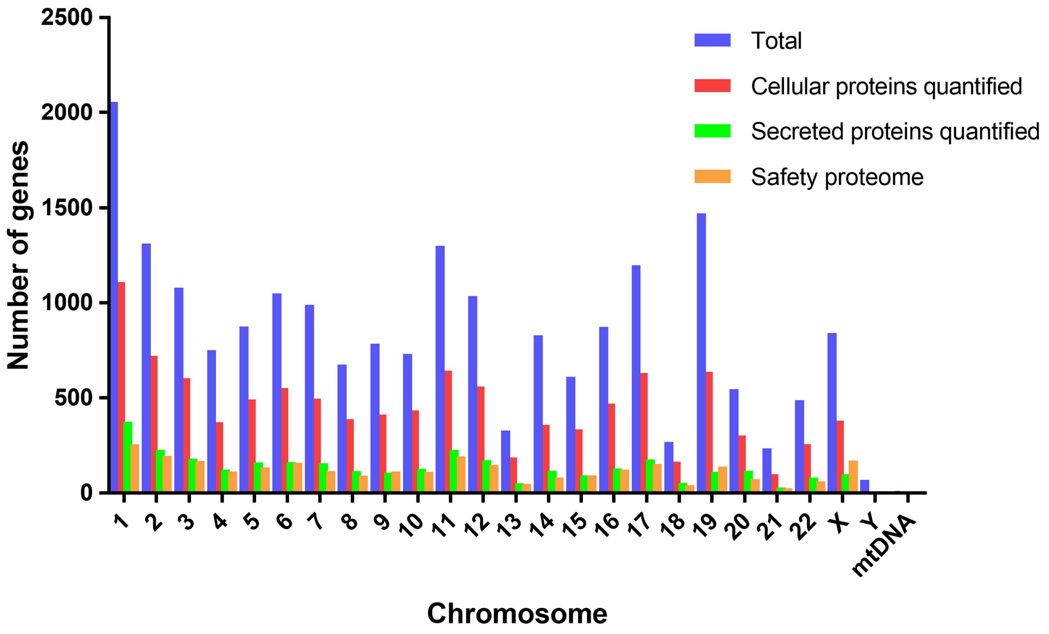

**Supplementary Figure 3**

Composition of the secretomes (quantified proteins). Venn diagram (A) and pie chart (B) of secretion pathways.

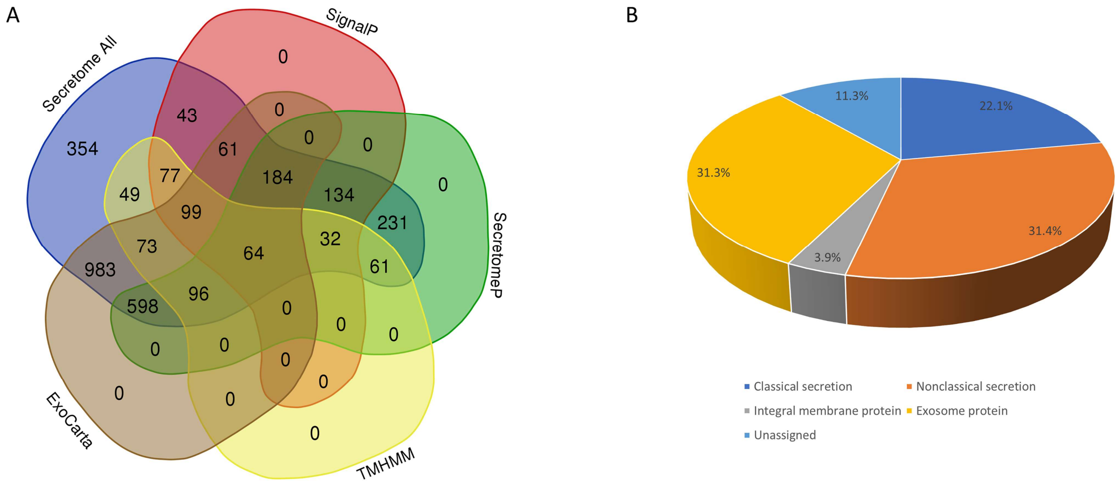

**Supplementary Figure 4**

Distribution of protein intensity (LFQ) from profiling experiments of cellular proteome (A) or secretome (B).

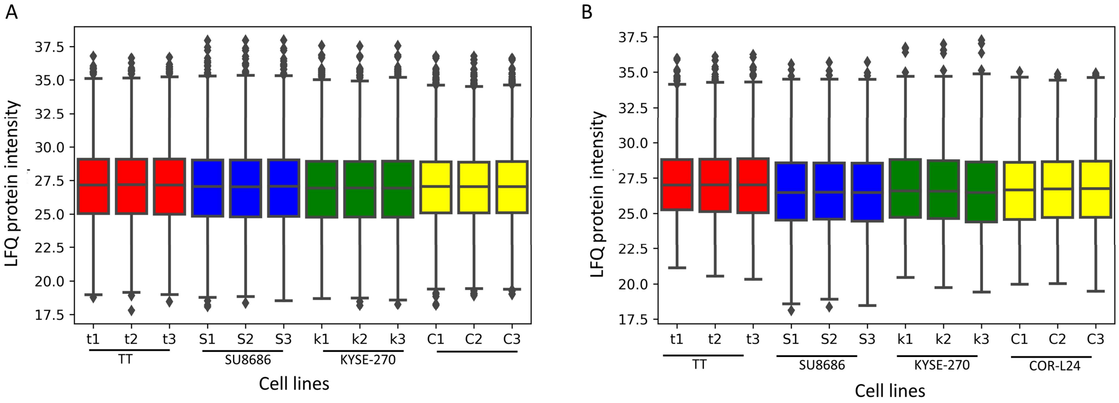


**Cellular proteome**

**Secretome**

**Supplementary Figure 5**
Dynamic range of the composite cell line cellular proteome estimated by iBAQ.

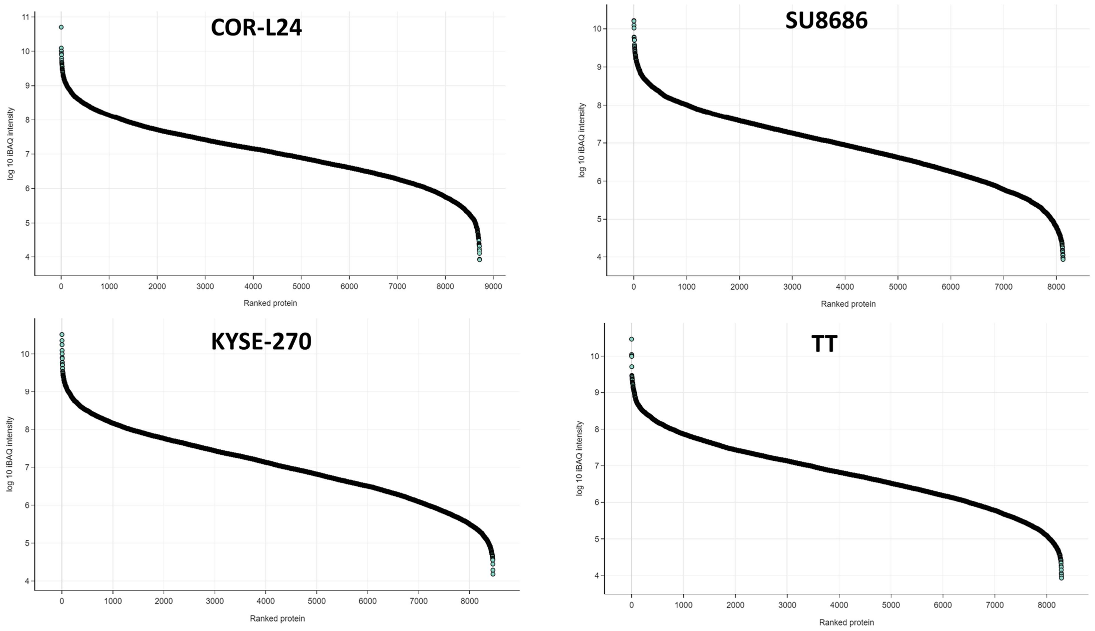

**Supplementary Figure 6**

Dynamic range of the composite cell line secretome estimated by iBAQ.

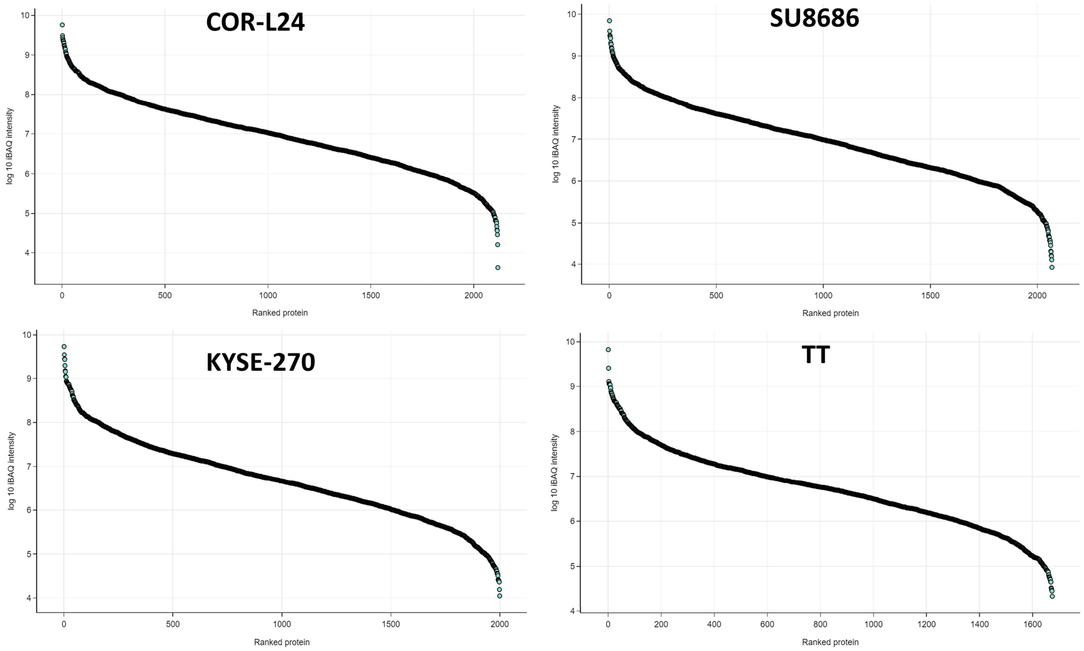

### Supplementary Figure 7

Distribution of the median logarithmic protein intensity (iBAQ) of all proteins quantified in cellular proteome (A) or secretome (B).

A

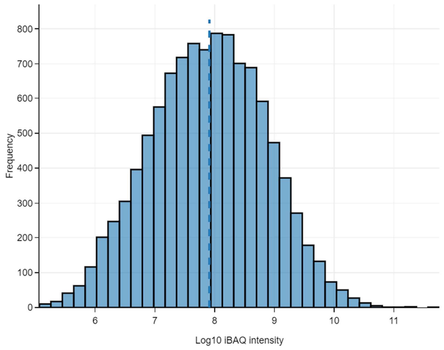


**Cellular proteome**

B

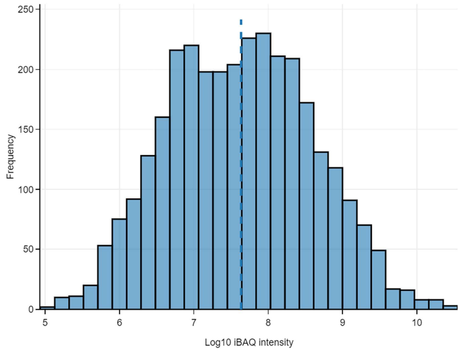


**Secretome**

**Supplementary Figure 8**

Matrix representation of scatter plots and Pearson correlation values of the label-free protein abundances of triplicates of each cell line cellular proteome(A) and secretome (B) against the others. Missing values were imputed based on a normal distribution (width = 0.3, down-shift =1.8)

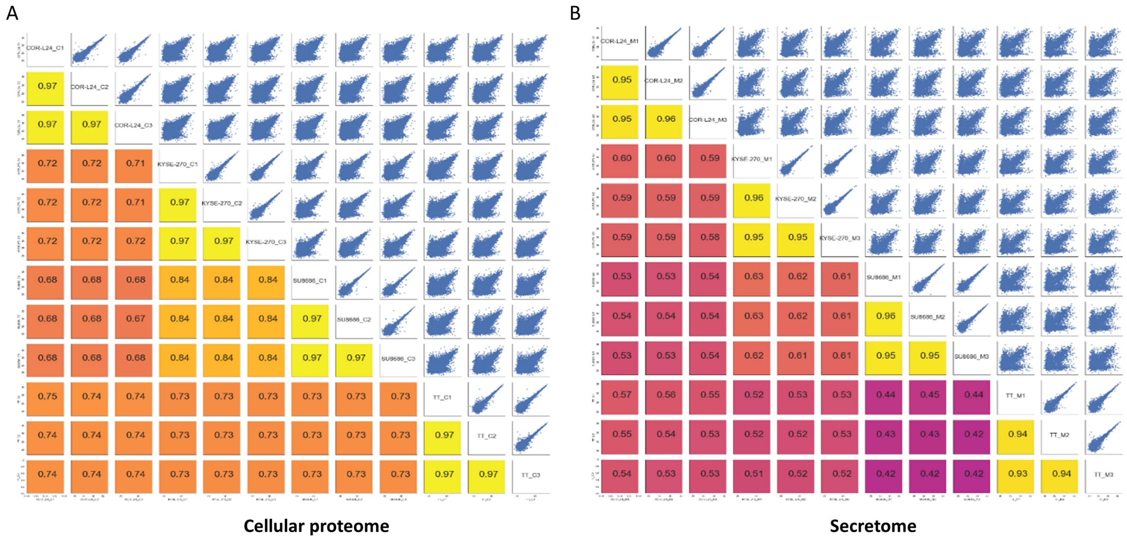

### Supplementary Figure 9

PCA and unsupervised hierarchical clustering based on label-free proteome quantification in cellular proteome (A and C) and secretome (B and D).

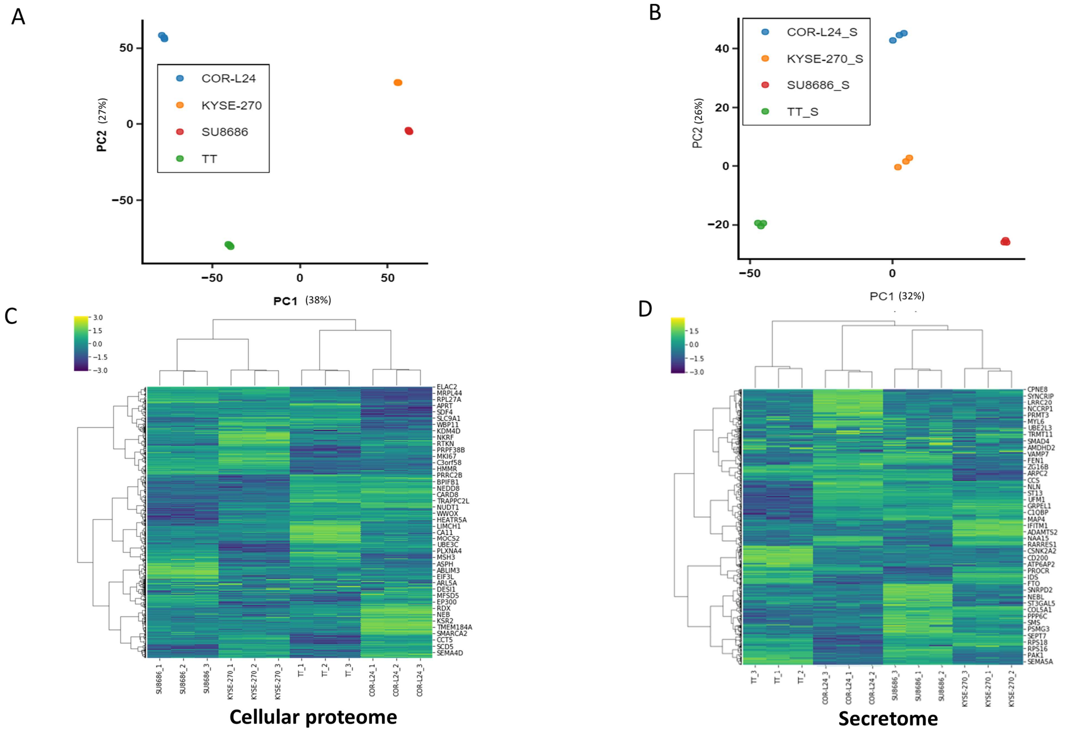

**Supplementary Figure 10**

Safety-related genes across organ classes.

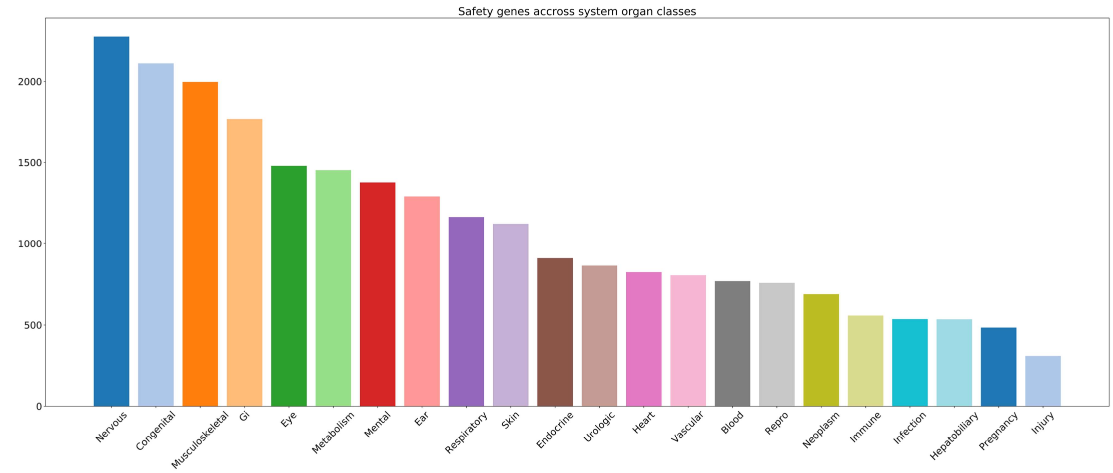
